# Dual-plane 3-photon microscopy with remote focusing

**DOI:** 10.1101/687202

**Authors:** Kevin T. Takasaki, Dmitri Tsyboulski, Jack Waters

**Affiliations:** Allen Institute for Brain Science, 615 Westlake Ave N, Seattle, WA 98109, USA; Janelia Research Campus, Howard Hughes Medical Institute, Ashburn, VA 20147, USA

## Abstract

3-photon excitation enables *in vivo* fluorescence microscopy deep in densely labeled and highly scattering samples. To date, 3-photon excitation has been restricted to scanning a single focus, limiting the speed of volume acquisition. Here, for the first time to our knowledge, we implemented and characterized dual-plane 3-photon microscopy with temporal multiplexing and remote focusing, and performed simultaneous *in vivo* calcium imaging of two planes deep in the cortex of a pan-excitatory GCaMP6s transgenic mouse. This method is a straightforward and generalizable modification to single-focus 3PE systems, doubling the rate of volume (column) imaging with off-the-shelf components and minimal technical constraints.

## 1. Introduction

Multiphoton laser-scanning fluorescence microscopy has enabled numerous studies of the structure and function of neural circuits *in vivo*. To date, the majority of these studies have been performed with two-photon excitation (2PE) in which the energy of two near-infrared photons is simultaneously absorbed to generate fluorescence at visible wavelengths (1, 2). The quadratic dependence of excitation on intensity provides optical sectioning and background reduction in the presence of dense fluorescent staining and volume scattering of excitation light. If scattering is sufficiently strong, however, such as when attempting to image directly through intact bone (3) or at many scattering lengths deep in brain tissue (4, 5), the second-order nonlinearity becomes insufficient to suppress fluorescence generated by out-of-focus excitation light, inevitably obscuring the in-focus signal. In such cases, three-photon excitation (3PE) microscopy has been shown to penetrate further and enable high-resolution fluorescence imaging beyond the depth limit of 2PE, enabling for example imaging directly through the skull (6) and into subcortical layers (7). The longer, short-wave IR wavelengths typically used and third-order nonlinearity in 3PE extend the scattering length and further reduce out-of-focus excitation, respectively.

A significant limitation of 3PE microscopy is the slow acquisition speed which can be over an order of magnitude slower than 2PE imaging. State-of-the-art multiplane and multi-region 2PE methods can acquire up to 10 standard raster-scanned planes at 10 Hz volume rate (8–10). At present, state-of-the-art 3PE imaging has been demonstrated with an acquisition rate equivalent to a single 2PE plane at roughly 10 Hz (11–13). Two factors contributing to the comparatively slow acquisition of 3PE are first, the low laser repetition rates used thus far for 3PE imaging, and second, the early stage of technical development. At present, the pulse repetition rates used in 3PE microscopy have been typically 1 MHz or less, far below the standard 80 MHz repetition rate in 2PE, limiting the acquisition speed to at most 15 Hz for 256×256 images sampled with 1 pulse per pixel. Recent work, however, indicates 3PE at repetition rates above 1 MHz is practical and might be leveraged for faster imaging. Experiments with a tunable repetition rate source for 2PE determined an ideal range of 1 to 10 MHz for deep 2PE imaging (14), and a recent study by Yildirim et al. (12) postulated a maximum safe repetition rate of 10 MHz for 3PE which, if implemented, could dramatically increase the acquisition speed. Though theoretically promising, whether 3PE at pulse rates above 1 MHz is beneficial for imaging and how to leverage additional pulses toward increasing acquisition speed still remain to be investigated.

To address the challenge of utilizing higher repetition rates, adapting volume imaging techniques from 2PE microscopy to 3PE is clearly sensible. A recent success has been demonstrating extended depth-of-focus scanning with Bessel beam 3PE (15, 16), but 2PE methods for imaging multiple planes simultaneously (9, 17–19) have not yet been adopted. Here, our goal was to implement 3PE multiplane imaging to accelerate volume acquisition without reducing the single plane framerate, as well as to minimize the necessary increase in excitation power and sample heating by optimizing excitation efficiency.

## 2. Multiplane Imaging

Compared with 2PE multiplane imaging methods, there are important considerations for implementing multiplane imaging with 3PE. Pulses generated by commercial OPA’s for 3PE are generally shorter than those typically used in 2PE imaging (~50 fs vs. ~200 fs transform-limit). Because of the larger spectral bandwidth, focus-shifting methods based on varying refractive index such as commonly used electro-optic and acousto-optic lenses, as well as liquid crystal spatial light modulators, are complicated by the need to characterize and limit chromatic aberration. Thus, recent work in correcting and shifting the 3PE focus with adaptive optics and remote focusing have relied on mirror-based approaches (20, 21). Moreover, because of the higher-order nonlinearity, signal generation by 3PE is more sensitive to the size of the focal volume. For 3PE, the rate of fluorescence emission at focus can be calculated to increase proportionally with the NA^2^, therefore benefitting from maintaining high NA, diffraction-limited focusing more than 2PE which in sufficiently thick samples is independent of the excitation NA (7). Efficient excitation is particularly important in multiplexing 3PE because water absorption and tissue heating by 1300 nm light places limits on the average power in the sample and therefore the pixel rate and number of planes that can be imaged at a given depth. With these factors in mind, we resolved to avoid chromatic effects while maintaining a high NA of ~0.9 by shifting the excitation focus with a commercial, plan-achromat air objective in a remote-focusing configuration.

Here, we describe simultaneous 3P imaging of two planes displaced by up to ±50 μm axially each at 7 Hz framerate with a 2 MHz effective repetition rate. We designed and constructed a dual-plane 3PE laser-scanning microscope using principles of temporal (time-division) multiplexing (17) and remote focusing (22, 23) to separate the spatiotemporal intensity distributions of the two excitation foci. We took advantage of a recently demonstrated demultiplexing circuit developed for multiplane 2PE microscopy (19) to separate analog signals from the two planes into individual channels. To maintain optimal 3PE imaging and minimize tissue heating, we characterized and minimized pulse dispersion and degradation of the remote focus. Finally, we validated dual-plane 3PE *in vivo* calcium imaging deep in the cortex of an awake, headfixed transgenic mouse. To our knowledge, the description of dual-plane, multiplexed 3PE microscopy is novel. We also note that this method is a straightforward and generalizable modification to single-plane 3PE systems, doubling the rate of volume acquisition with off-the-shelf components and minimal technical constraints.

## 3. Experimental setup and procedures

### 3.1 Optical setup

A schematic of the dual-plane remote focusing 3P microscope is shown in Figure 1. Excitation laser pulses were generated by a commercial non-collinear OPA (Opera-F, Coherent) operating at 1300 nm (idler beam) pumped by a 40 W fiber laser (Monaco, Coherent) operating at 1040 nm. Excitation pulses at the output of the laser were measured to be nominally 2 μJ with a 50 fs pulse duration (PulseCheck, APE). The beam was attenuated by a half-wave plate (AHWP-1600, Thorlabs) and polarizing beam splitter (PBS104, Thorlabs), then directed into a single-prism (N-SF11, PS855, Thorlabs) pulse compressor for dispersion compensation. The beam exited the compressor at a lowered height and was picked off by a D-mirror. A second half-wave plate and polarizing beam splitter was used to divide the pre-chirped beam into two paths for either remote focusing or temporal delay. The transmitted beam path (remote focusing; path A) was expanded by a 1.5x beam expander (AC254-100-C; AC254-150-C, Thorlabs) then entered the microscope platform (MIMMS, Sutter Instruments) via a periscope. The reflected beam path (temporal delay; path B) was reflected by a series of mirrors over ~5 m to delay the pulses by roughly 15 ns relative to path A, then demagnified 2x by a beam reducer (BE-02-05-C, Thorlabs) to compensate for the long propagation distance before entering the microscope via a separate periscope, in the process becoming p-polarized, and recombining with the path A beam at another polarizing beam splitter.

**Figure 1.**
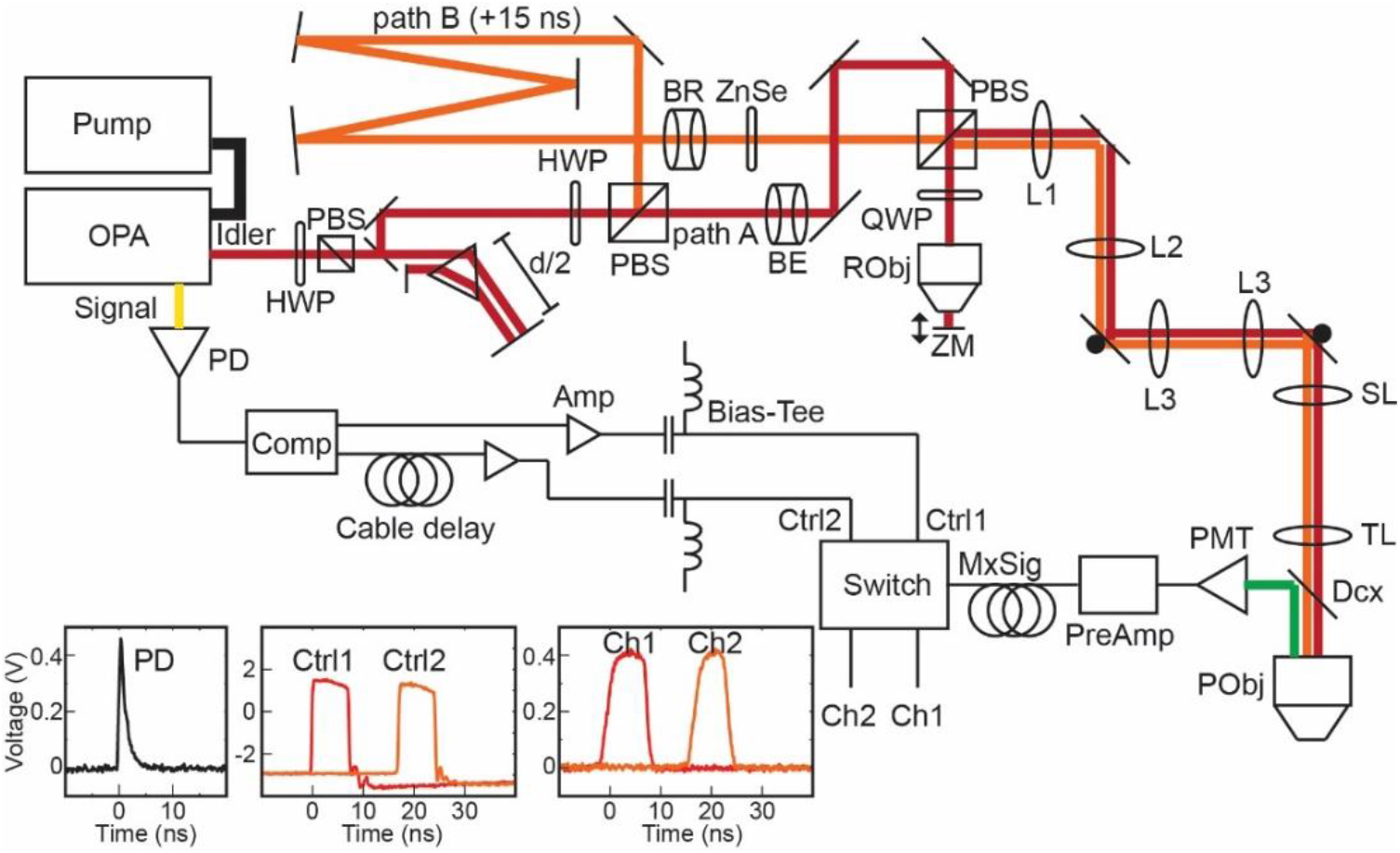
Schematic of Dual Plane 3P Microscope. System schematic of dual plane 3PE microscope with analog demultiplexing electronics. Abbreviations: OPA – optical parametric amplifier, HWP – half-wave plate, PBS – polarizing beam splitter, BE – beam expander (1.5x), BR – beam reducer (0.5x), QWP – quarter-wave plate, RObj – remote focus objective, ZM – z-translation mirror, L1 – doublet lens (f=150 mm), L2 – doublet lens (f=100 mm), L3 – Plössl lens (f = 50 mm), SL – scan lens (f=50 mm), TL – Plössl tube lens (f=200 mm), Dcx – dichroic beam splitter, PObj – primary objective, PMT – photomultiplier tube, PreAmp – pre-amplifer, MxSig – multiplexed signal, Amp – amplifier, Comp – comparator circuit, PD – photodiode. Inset: Oscilloscope traces of photodiode output (left), gate control pulses (center), and demultiplexed channel detection windows exhibited by switching a 0.5 VDC signal input (right).

Initially, the p-polarized path A beam passed through the recombining beam splitter. The linearly polarized beam was circularly polarized by a quarter-wave plate (AQWP-1600, Thorlabs), then entered the remote objective (LCPLN50XIR, 0.65 NA, Olympus). Filling the remote objective pupil allowed a maximum excitation NA of 0.65×1.33 ≈ 0.9 which was achieved by imaging the remote pupil onto the primary objective pupil at a magnification of 8/3. The focused beam was reflected back through the objective by a mirror mounted on a manual translation stage, and became s-polarized by reverse passage through the quarter-wave plate. The combined beams then entered a 4f-relay (AC254-150-C; AC254-100-C, Thorlabs) configured to image the back pupil of the remote objective onto the scanning galvanometers (6215H, Cambridge Technologies). The x- and y- galvos were also imaged onto each other by a pair of Plössl scanning lenses, then imaged onto the pupil of the primary objective (XLPLN25XWMP2, Olympus) by a scan lens (SL50-3P, Thorlabs) and Plössl tube lens. Emitted green fluorescence was separated from incoming excitation light by a dichroic beam splitter (FF735-Di02, Semrock), then filtered by a bandpass filter (ET525-70m-2p, Chroma), and detected by a GaAsP photomultiplier tube (H10770PA, Hamamatsu). Photocurrents from the PMT were amplified by a high-speed pre-amplifier (DHPCA-100, Femto, 200 MHz BW, 10^3^ V/A) and sent into the demultiplexing electronics.

### 3.2 Demultiplexing electronics

The signal beam at 855 nm was attenuated by a beamsplitter and neutral density filters, then sampled by an amplified photodiode (DET10A, Thorlabs) with a ~1 ns rise time. The photodiode pulse was sent into a comparator (LTC6957-HMS3, Analog Devices) which converted the input pulse waveform into two square pulse outputs of ~8 ns duration, one of which was delayed by 15 ns with cabling. The square pulses were scaled to a peak-to-peak amplitude of 5 V with an amplifier (GVA-83+, MiniCircuits) and shifted to a common mode of −1 V by a bias tee (ZFBT-4R2GW+, MiniCircuits). The conditioned pulses were used to gate an RF switch (CMD196C3, Custom MMIC) whose input signal was switched between two output terminals connected to channels 1 and 2 of the digitizer (see Fig.1 inset). The analog voltage signal from the pre-amplifier was delayed with cabling to temporally align with the channel windows before being input into the switch. Analog-to-digital conversion of the DAQ channels was also gated by 1 MHz pulses from the TTL sync output from the pump laser.

### 3.3 Third-order interferometric autocorrelation

The autocorrelator was constructed from a 50:50 non-polarizing beam splitter (BS015, Thorlabs) and a pair of mirrors, one of which was mounted on a piezo translator (nPFocus400, nPoint). Piezo movement was controlled in ScanImage (ScanImage Premium, Vidrio LLC) as a fast Z-axis scanner and synchronized with imaging of fluorescence from 50 μM fluorescein solution. Signal detected from fluorescein was averaged line-by-line and plotted against relative pulse delay calculated from the piezo scan position. Interferometric autocorrelation traces were numerically simulated in MATLAB (Mathworks) for Gaussian pulses of specified bandwidth and group dispersion delay (GDD) as chirp applied in the Fourier domain.

## 4. Results

Of significant concern with using remote focusing in 3PE is the additional pulse dispersion introduced by the remote focusing unit, particularly by the double pass through the remote objective. To compensate dispersion, the idler beam was passed through a single-prism compressor which allowed substantial pre-chirp with robust tunability and a compact footprint. Because this pre-chirp was applied to both the remote focusing and temporal delay paths, it was critical to ensure the pulses in the two paths encountered identical GDD and could be ideally compensated at the microscope focus by the same compressor configuration. To characterize the pulse at the microscope focus, we constructed a Michelson interferometer in the beam path and measured 3PE-generated fluorescence in fluorescein while modulating the path length of one interferometer arm with a piezo-actuated mirror (Fig. 2a). In our setup, we used a commercial piezo scanner optimized for millisecond displacements of a microscope objective over travel distances of up to half a millimeter, but identical measurements could be obtained with a less specialized actuator and implemented at low cost compared with a commercial autocorrelator. Green fluorescence detected from 50 μM fluorescein excited at 1300 nm displayed a power dependence consistent with a cubic nonlinearity (n = 3.04 ± 0.07; Fig. 2b). Delaying one interferometer arm resulted in fringes in the fluorescence level reflecting the third-order interferometric autocorrelation (TIAC) (24, 25) of the excitation pulse. We numerically generated TIAC traces for pulses of varying widths and chirps, and determined the best fit of the empirical data by the least-squares error. To cross-validate our measurement of the pulse width, we measured the pulse profile with minimal optics in the path, removing all but the prism compressor, beam expander, and objective, and tuned the compressor length to minimize the width of the TIAC. With the compressor set to a distance of 44 cm prism separation (−4500 fs^2^ GDD), the measured trace was best fit by the simulated TIAC of a Gaussian pulse with intensity full-width at half-maximum (FWHM) of 48 fs. This was consistent with estimates of the transform-limited FWHM of the pulse obtained from both measurements of the spectral bandwidth by spectrometer and of the autocorrelation of the undispersed, free-space beam with a commercial autocorrelator (PulseCheck, APE). We similarly optimized the pulse through the full remote focusing path (path A) and obtained a minimal pulse width of 53 fs with 92 cm prism separation, corresponding to ~11000 fs^2^ GDD (Fig. 2c). Subsequently, measuring the temporally delayed path (path B) with 92 cm prism separation, the FWHM of the chirped pulse was ~140 fs at focus with a relative negative GDD of −2400 fs^2^ which we compensated by inserting a disk of ZnSe of 5 mm thickness (Fig. 2d). To validate these measurements and examine the effect on excitation efficiency, we removed the interferometer and imaged a fluorescent bead sample with and without the ZnSe disc. We observed nearly a 10-fold difference in fluorescence for equal excitation energy (Fig. 2e), demonstrating the importance of proper dispersion compensation for limiting excitation power and tissue heating.

**Figure 2.**
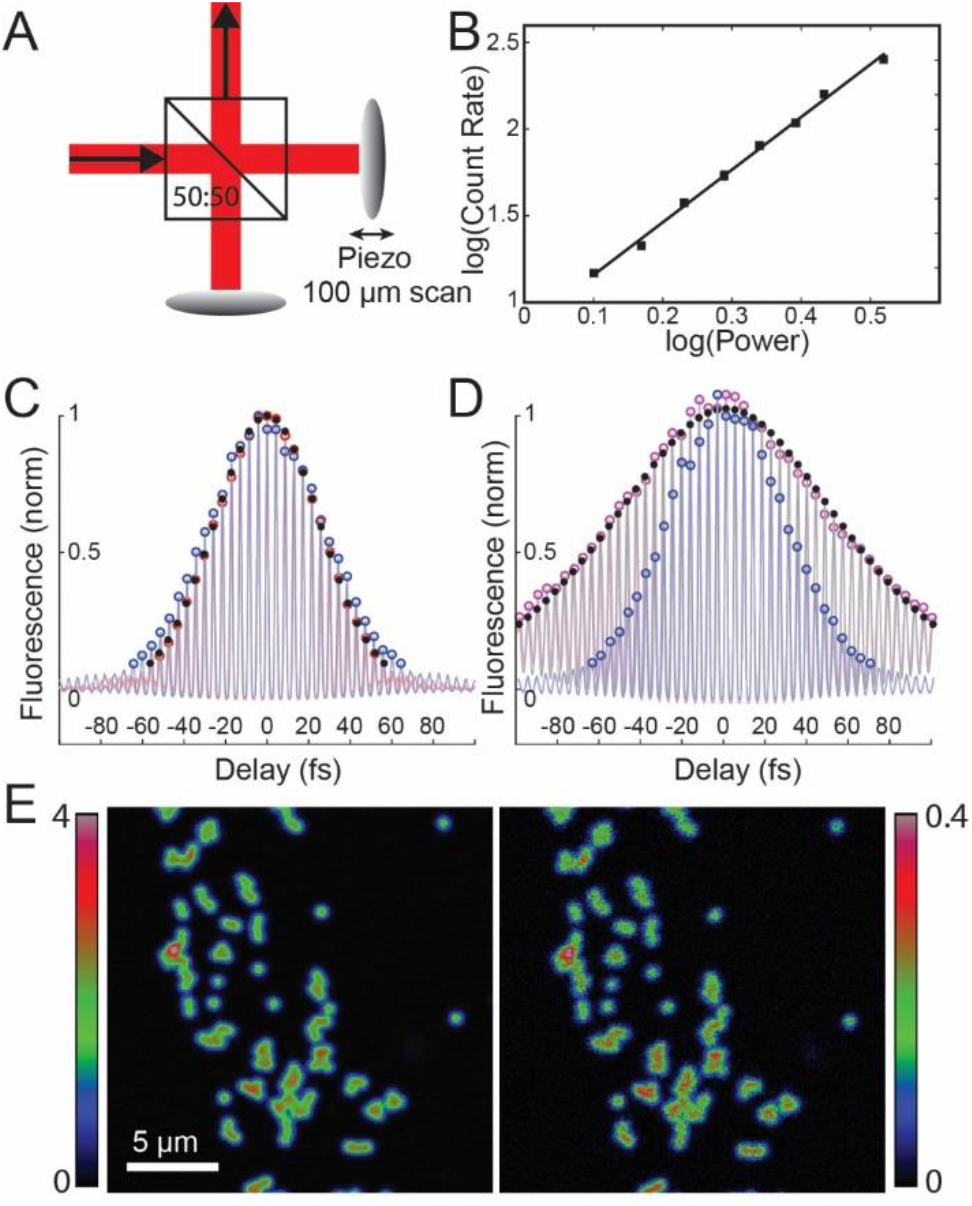
Pulse dispersion characterization and compensation. **A**, illustration of interferometric autocorrelator setup inserted into the beam path. A 50:50 beam splitter splits incoming pulses between two arms of nearly equal length which are reflected back by mirrors, one of which is mounted on a piezo-actuated translation stage. The pulses are recombined leading to interference and amplitude modulation of the excitation focus. **B**, log-log plot of fluorescence vs. excitation power in 50 μM fluorescein with a linear fit of slope n = 3.04. **C**, plot of peak-normalized fluorescence detected in fluorescein vs. inter-pulse delay calculated from position of interferometer mirror for minimal optics (d = 44 cm; red circles) and path A (d = 92 cm; blue circles) compared with the minimal best-fit from simulation (FWHM = 48 fs; black dots). **D**, plot of peak-normalized fluorescence for path B without the ZnSe disc (d = 92 cm; purple circles) and with the ZnSe disc (5 mm ZnSe, d = 92 cm; blue circles) compared with the best-fit from simulation with negative chirp (GDD = −2400 fs^2^; black dots). **E**, images of 500 nm fluorescent beads in water imaged with (left panel) and without (right panel) the ZnSe disc. Color scale: mean photon rate of 0 – 4 counts/pixel (left) and 0 – 0.4 counts/pixel (right, 10x gain).

We next examined the range of diffraction-limited remote focusing. Principles of remote focusing can be found detailed in the literature (23, 26), and here we provide only a summary description. Briefly, our aim was to modulate the wavefront in the primary objective back pupil to cause axial repositioning of the focus without distorting its shape, i.e. while maintaining the resolution in the native focal plane. In the remote focusing approach, the modulated wavefront is produced in the back pupil of the remote objective and imaged onto the pupil of the primary objective, and so the range of diffraction-limited refocusing is constrained by the axial range over which the wavefront generated remotely corresponds to the ideal wavefront required for proper focusing. One consequence of this correspondence requirement is that the axial range typically shrinks with increasing NA, as the contribution of higher-order spherical terms, which are more difficult to correct precisely, grows with both the NA and magnitude of axial displacement. Thus, the axial range of high NA refocusing may be determined not only by the specifications of the remote and primary objectives, but also their optical design quality and tolerances. Given these considerations, we began by measuring the axial range of our remote focusing system empirically.

The PSF measured with 500 nm beads at the native focal plane, i.e. with the remote mirror positioned at the remote focal plane, exhibited a FWHM of 0.67 μm laterally and 1.9 μm axially. By translating the remote mirror, we observed axial displacement of the imaging plane up to ±50 μm while maintaining near diffraction-limited resolution (Fig. 3a). Beyond 50 μm, the focal volume progressively enlarged with significantly reduced resolution (Fig. 3b). The cubic nonlinearity implies that even the common definition of a Strehl ratio threshold of 0.8 for diffraction-limited performance (Maréchal criterion) will lead to a 50% reduction in fluorescence from a point emitter, approximately the signal reduction we observed at ±50 μm displacement. Indeed, the PSF measured at ±50 μm was only slightly enlarged relative to its size at the native imaging plane, closely matching theoretical predictions for the size of a diffraction-limited focus for an NA of 0.9 (Fig. 3c-d). Based on these measurements, we prescribed the range of the remote-focusing plane to within ±50 μm.

**Figure 3.**
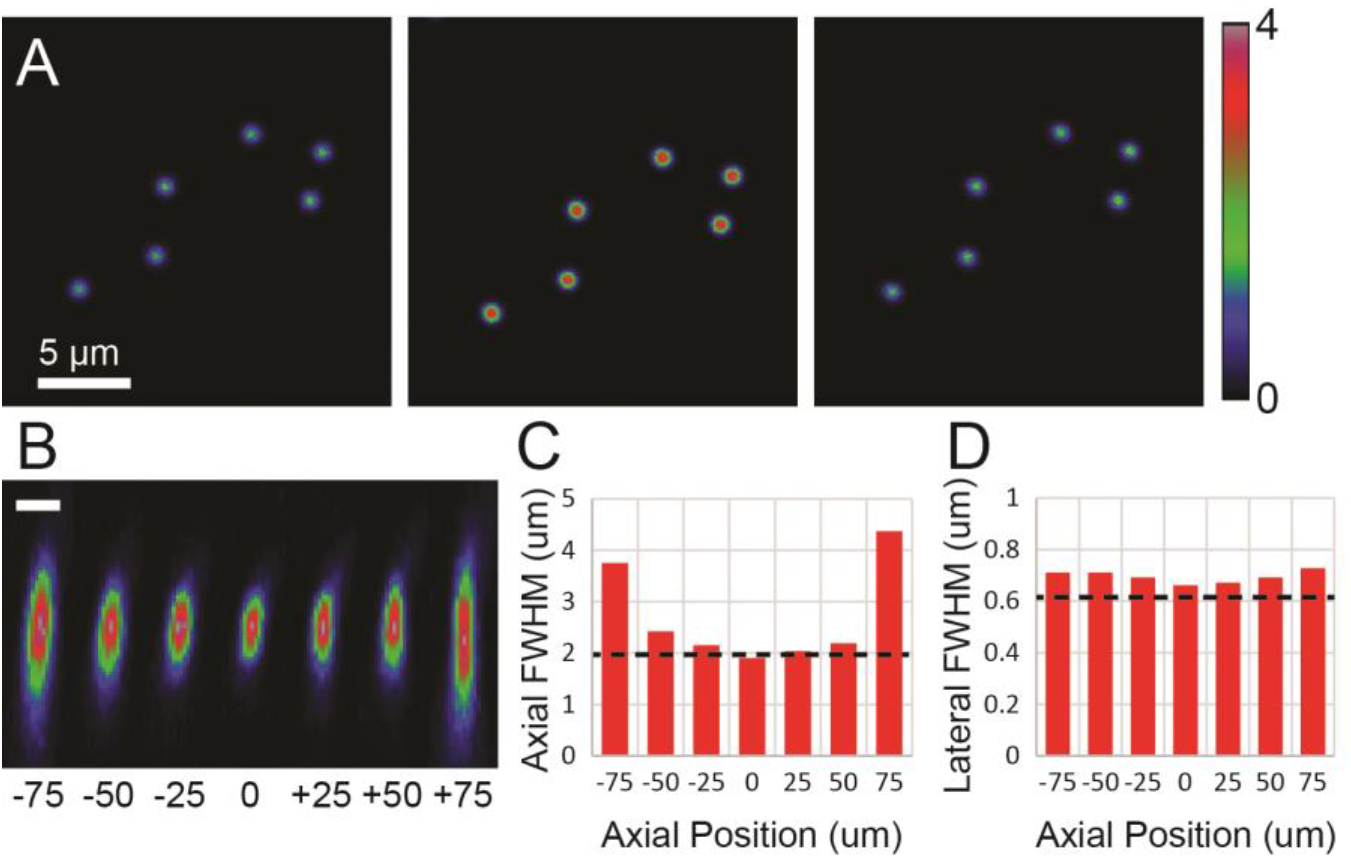
Characterization of PSF with remote focusing. **A**, images of 500 nm fluorescent beads taken in the remote focusing plane for axial displacements of −50 μm (left), 0 μm (center), and +50 μm (right) with equal average power. Color scale: Mean photon rate 0 – 4 counts/pixel. **B**, xz-projection of bead images. Scale bar: 1 μm. **C**, graph of measured axial FWHM for different remote focused plane positions. Dashed line denotes theoretical axial FWHM for NA = 0.9 focus. **D**, graph of measured lateral FWHM for plane positions as in C. Dashed line denotes theoretical lateral FWHM for NA = 0.9 focus convolved with a 0.5 μm spherical shell representing the bead sample.

Finally, we performed dual-plane calcium imaging of neurons deep in the cortex of a transgenic mouse expressing the genetically encoded calcium indicator GCaMP6s (27) in excitatory neurons throughout cortex (Slc17a7-IRES-Cre;Ai94). We simultaneously imaged two planes located at 600 and 650 μm deep, beyond the 2PE depth limit in these transgenic mice (5), in two 10 minute sessions. Between sessions, we exchanged the planes between path A and path B excitation to compare remote focusing at both ±50 μm planes with conventional focusing at the native imaging plane (Fig. 4a-b). Consistent with our measurements of the PSF, signal reduction in the planes imaged at ±50 μm were compensated by modest changes in illumination power (15 mW path B vs. 17 mW path A at 650 μm). Image quality appeared identical between remote and normal focusing with similar features appearing in both movies, including transient activity in small puncta and dendrites. We compared fluorescence traces from spontaneously active neurons after motion correction and segmentation with Suite2p (28) and observed calcium transients of similar amplitude, frequency, and time course in extracted fluorescence traces (Fig. 4c-d).

**Figure 4.**
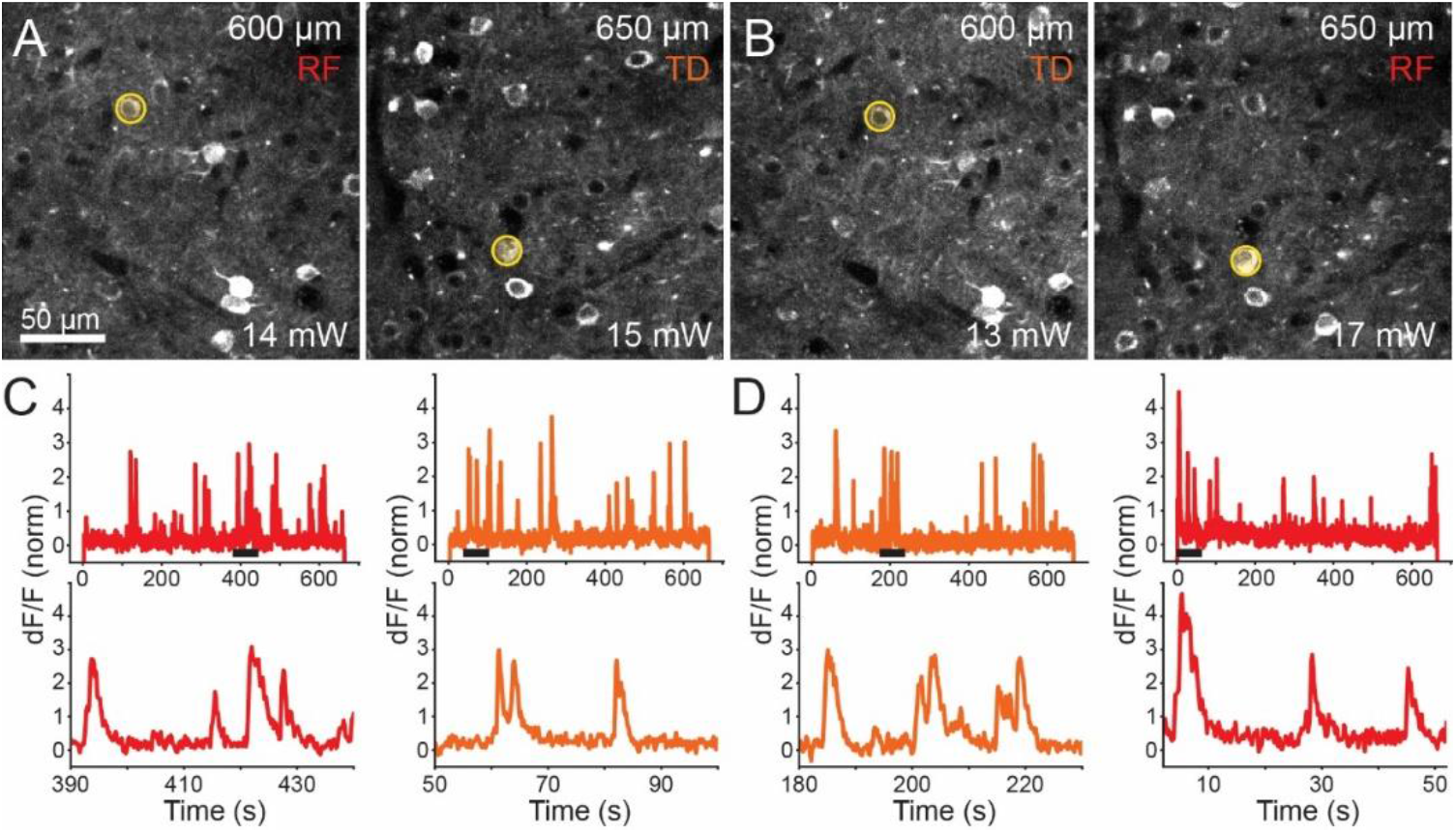
Dual plane calcium imaging of neurons deep in cortex. **A**, maximum intensity projections of motion corrected movies from simultaneously acquired planes located at z = 600 μm (left) and z = 650 μm (right) below the cortical surface imaged with the remote focus (RF, path A) and temporally delayed conventional focus (TD, path B), respectively. **B**, maximum intensity projections of motion corrected, simultaneously acquired movies from planes in A with the remote focus and conventional focus exchanged. **C**, dF/F normalized fluorescence traces extracted from neurons highlighted in A from movies acquired 50 μm above (red) and at (orange) the native imaging plane. Top panel: dF/F trace over the entire movie. Bottom panel: underlined section of full trace. **D**, dF/F normalized fluorescence traces extracted from neurons highlighted in B from movies acquired at (orange) and 50 μm below (red) the native imaging plane. Top panel: dF/F trace over the entire movie. Bottom panel: underlined section of full trace.

## 5. Discussion

The use of 3PE microscopy in neuroscience has grown considerably in recent years, driven by applications requiring high-resolution imaging through strongly scattering and densely fluorescent samples. However, data acquisition speeds with 3PE at present are an order of magnitude slower than comparable acquisition with 2PE, generally increasing the experimental burden and therefore constraining the practicality of using 3PE in situations where it is technically needed. In this work, we performed 3PE imaging in deep cortex at 2 MHz with total average power (30 mW) well below estimates of the thermal limit (6), indicating the possibility of further increasing both the pulse repetition rate and imaging speed. The low average power required per plane was made possible by controlling the spatiotemporal profile of excitation light to minimize the pulse width and size of the focal volume so that they remained close to their physical limits. We implemented dual plane imaging using techniques taken directly from 2PE microscopy with consideration given to the different characteristics of 3PE and anticipate further leveraging of established methods to be helpful, as has also been demonstrated with focus shaping and adaptive optics.

Several improvements to this work are foreseeable. The first challenge is to extend the number and range of imaging planes. Here, we did not move the remote mirror during the acquisition as is typically done in multiplane 2PE systems. The current frame rate of ~7 Hz is borderline for sequential plane repositioning, but increasing the pulse repetition rate and frame rate with beam multiplexed and/or resonant scanning should permit fast translations of the remote focusing planes with a reasonable overall volume acquisition rate. The range of remote focusing is currently limited by the remote objective. We selected the remote objective used here with the goal of maintaining high NA, diffraction-limited focus quality and high transmission at 1300 nm with an off-the-shelf part, albeit with an acceptable limit on the range of refocusing. In theory, diffraction-limited, high-resolution imaging over hundreds of microns axially should be achievable if a more ideal objective (e.g. 20X, 0.8 NA, IR air objective) were acquired or designed. Another critical challenge is to limit the average power and sample heating as the number of planes is increased. Here, gains from more efficient excitation with adaptive optics (20, 29), brighter indicators (30), and sample-tailored illumination (31) would contribute to reducing the average illumination power per plane.

## 6. Conclusion

We have described the implementation and characterization of a dual-plane 3PE laser-scanning microscope with temporal multiplexing and remote focusing, the first description of such to our knowledge. The method is a straightforward and low-cost modification of single-focus 3PE microscopes, and is readily applicable to *in vivo* imaging.

## Acknowledgments

We wish to thank Daniel Flickinger and Natalia Orlova for their expertise and helpful discussions. We wish to thank the founder of the Allen Institute for Brain Science, Paul G. Allen, for his vision, encouragement, and support.

